# Understanding biases in ribosome profiling experiments reveals signatures of translation dynamics in yeast

**DOI:** 10.1101/027938

**Authors:** Jeffrey A. Hussmann, Stephanie Patchett, Arlen Johnson, Sara Sawyer, William H. Press

## Abstract

Ribosome profiling produces snapshots of the locations of actively translating ribosomes on messenger RNAs. These snapshots can be used to make inferences about translation dynamics. Recent ribosome profiling studies in yeast, however, have reached contradictory conclusions regarding the average translation rate of each codon. Some experiments have used cycloheximide (CHX) to stabilize ribosomes before measuring their positions, and these studies all counterintuitively report a weak negative correlation between the translation rate of a codon and the abundance of its cognate tRNA. In contrast, some experiments performed without CHX report strong positive correlations. To explain this contradiction, we identify unexpected patterns in ribosome density downstream of each type of codon in experiments that use CHX. These patterns are evidence that elongation continues to occur in the presence of CHX but with dramatically altered codon-specific elongation rates. The measured positions of ribosomes in these experiments therefore do not reflect the amounts of time ribosomes spend at each position in vivo. These results suggest that conclusions from experiments in yeast using CHX may need reexamination. In particular, we show that in all such experiments, codons decoded by less abundant tRNAs were in fact being translated more slowly before the addition of CHX disrupted these dynamics.

## Introduction

Translation is the process by which the assembly of a protein is directed by the sequence of codons in a messenger RNA. Ribosomes mediate this conversion of information from codons into amino acids through the sequential binding of tRNAs [42]. During the incorporation of each successive amino acid, there are several stages at which the identity of the codon being translated may potentially influence the speed with which a ribosome advances along a coding sequence. When a codon is presented in the A-site of a ribosome, an appropriate tRNA must diffuse into the A-site and successfully form a codon-anticodon base pairing interaction [15, 25]. tRNAs decoding different codons are expressed at different abundances [9, 43], suggesting that ribosomes could spend longer waiting for less abundant tRNAs to arrive [44]. Because translation is accomplished with fewer tRNA identities than there are codon identities, some codon-anticodon interactions involve non-Watson-Crick base-pairings [32, 16]. These so-called wobble pairings are thought to modulate the speed of decoding [24, 28, 41, 47]. Once a tRNA has arrived and base-paired, the speed of peptide bond formation between the C-terminal amino acid in the nascent chain and an incoming amino acid may be influenced by chemical properties of these amino acids [2]. The relative contributions of these effects to overall rates of translation remain poorly understood.

Because the genetic code that governs the process of translation maps 61 codon identities to only 20 standard amino acids, multiple synonymous codons can be used to encode most amino acids. There is a rich body of theoretical work on the role of translation speed as a selective force shaping synonymous codon usage [33], but the ability to directly measure the speed with which each codon is translated *in vivo* in order to test these theories has historically been lacking. The recent development of ribosome profiling, the massively parallel sequencing of footprints that actively translating ribosomes protect from nuclease digestion on messenger RNAs [21, 18, 20, 19], presents exciting opportunities to close this gap. Ideally, the millions of sequencing reads produced by a ribosome profiling experiment are snapshots of translation representing samples drawn from the steady state distribution of ribosomes across all coding sequences. The statistical properties of these snapshots can in theory be used to measure the relative speed with which each codon position is translated: the more often ribosomes are observed at a position, the longer ribosomes are inferred to spend at that position.

In practice, ribosome profiling studies in *Saccharomyces cerevisiae* using different experimental protocols have reached contradictory conclusions about the average decoding times of codon identities. Because yeast rapidly regulate translation when stressed and ribosomes cannot be instantaneously harvested from cells, the original ribosome profiling protocol of Ingolia et al. [21] pretreats cells with cycloheximide (CHX) for several minutes to stabilize ribosomes in place before the harvesting process begins. CHX is a small-molecule translation inhibitor that has been a staple of experimental approaches to the study of translation for decades. The exact mechanism of this inhibition, however, is not completely understood, with recent studies suggesting that CHX binds to a ribosome’s E-site along with a deacylated tRNA to block further translocation [38, 8]. The majority of the rapidly growing body of ribosome profiling experiments in yeast have followed this original CHX-pretreatment protocol [4, 14, 49, 11, 3, 2, 27, 29, 1, 30, 39]. Several groups have applied a variety of conceptually similar computational methods to the data produced by these experiments to infer the average speed with which each codon identity is translated. Counterintuitively, these groups have found that, on the whole, codons decoded by rare tRNAs appear to be translated faster than those decoded by more abundant tRNAs [36, 5]. Different theories have been advanced to explain these unexpected results, hypothesizing that the measured elongation rates reflect a co-evolved balance between codon usage and tRNA abundances [36], or that translation dynamics are dominated by interactions involving the nascent chain rather than the actual decoding process [5].

An alternative experimental protocol uses an optimized harvesting and flash-freezing process that allows CHX pretreatment to be omitted [21, 20, 27, 12, 13, 17, 34, 30, 48, 45]. Experiments performed with this protocol have revealed that treatment with CHX affects several high-level characteristics of footprinting data, including the distribution of lengths of nuclease-protected fragments in mammalian cells [22] and the amount of enrichment in ribosome density at the 5’ end of coding sequences in yeast [13]. In contrast to data produced using CHX pretreatment, several studies using this alternative protocol have reported that non-optimal codons are in fact translated more slowly [12, 45]. The source of this discrepancy between the statistical properties of measured ribosome positions with and without CHX pretreatment has been unclear, leading to uncertainty as to which measurements correspond to actual properties of *in vivo* translation dynamics.

Here, we present analysis of data from a large body of ribosome profiling studies in yeast to resolve these contradictory results. We find consistent differences between experiments performed with and without CHX pretreatment in how often ribosomes are measured with specific codon identities positioned in their tRNA binding sites. We also find unexpected patterns in how often ribosomes are found downstream of specific codon identities in experiments using CHX pretreatment. Together, these observations suggest that translation elongation continues for many cycles after the introduction of CHX, but that the amount of time ribosomes spend translating each codon under these perturbed conditions is quite different from the unperturbed dynamics.

## Results

### Pretreatment with cycloheximide consistently changes enrichments of codon identities at ribosomal tRNA binding sites

To characterize differences in the measured positions of ribosomes when cells are pretreated with CHX and when they are not, we compared the relative amount of time ribosomes spend at each codon position in data from many different ribosome profiling experiments. For each experiment, we mapped footprint sequencing reads to yeast coding sequences and assigned each read to the codon position in the A-site of the associated ribosome (the sixth codon from the 5’ end in a canonical 28nt footprint [21]). In each coding sequence, we divided the raw count of reads at each codon by the average across the coding sequence to produce a relative enrichment value for each position. To measure the average relative amount of time a codon identity spends in the A-site of a ribosome each time it is translated, we computed the mean of these relative enrichment values at all occurrences of each codon across all yeast coding sequences (Figure 1A). If the mean relative enrichment of a codon is higher than 1, translation of the codon is inferred to be slower than the average speed of its surroundings. Conversely, a mean relative enrichment value lower than 1 indicates that translation of a codon is faster than its surroundings.

**Figure 1:**
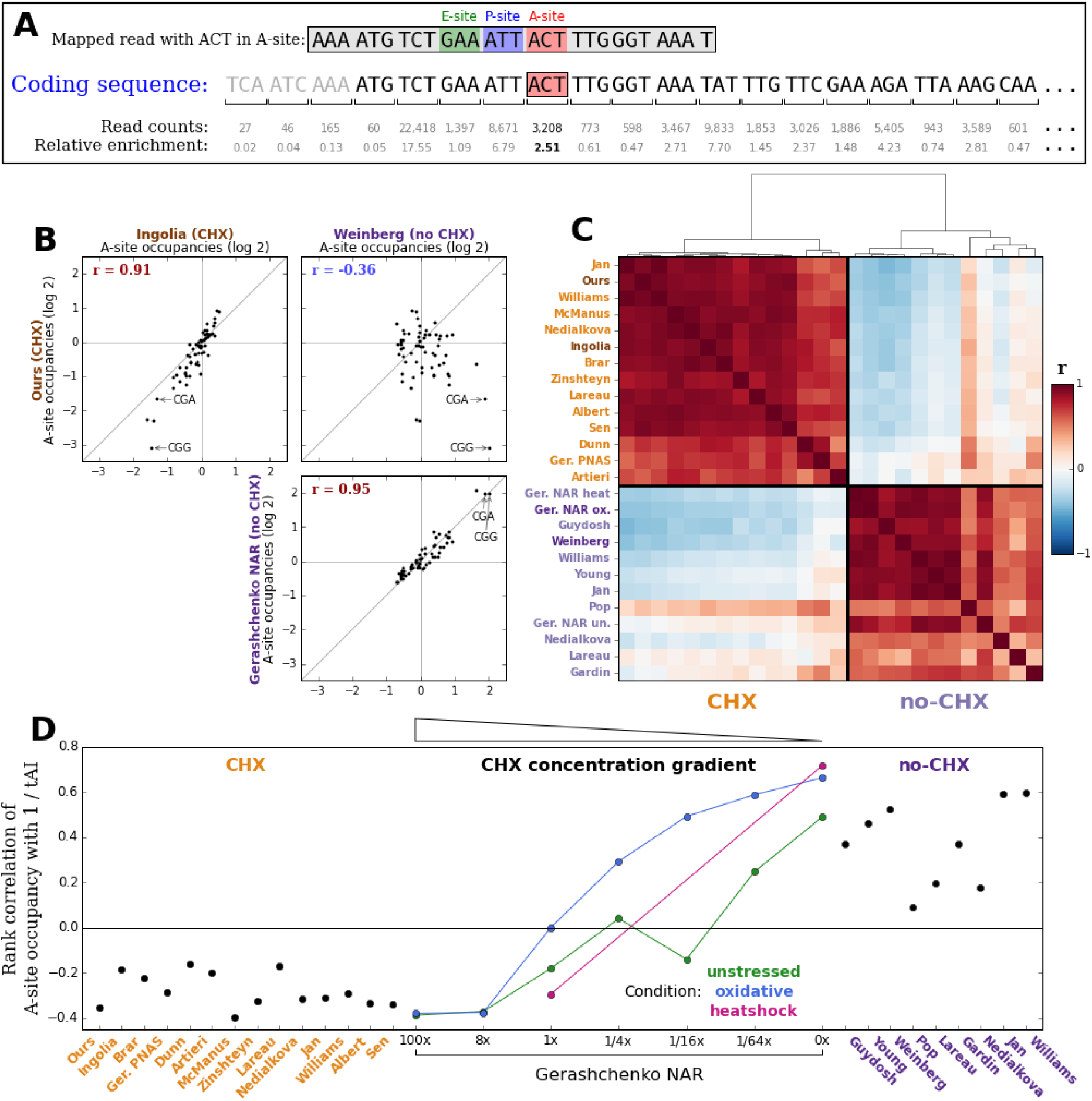
Experiments with and without CHX pretreatment report different A-site occupancies. **(A)** To measure how long a ribosome spends on average with each codon identity in its A-site, footprint sequencing reads (such as the boxed sequence on the top line) are mapped to yeast coding sequences and assigned to the codon position that the A-site of the ribosome was positioned over (red box in coding sequence). For each coding sequence, read counts at each position (shown below each codon) are divided by the average count across the coding sequence to produce relative enrichment values (shown on the bottom line). The average A-site occupancy of each codon identity (in this example, ACT) is then computed by averaging the relative enrichment at all occurrences of the codon (such as the bolded value) across all coding sequences. **(B)** Scatter plots comparing average A-site occupancies of all 61 non-stop codons between different pairs of experiments. A pair of experiments using CHX pretreatment (upper left) and a pair of experiments done without CHX pretreatment (lower right) report A-site occupancies with strong positive Pearson correlations, but these two internally consistent sets of values are strikingly different from each other (upper right). **(C)** Pearson correlations of average A-site occupancies of all 61 non-stop codons between representative experiments from many different studies in yeast, grouped by hierarchical clustering. Red indicates positive correlation and blue indicates negative correlation. Clustering separates experiments using CHX pretreatment (labeled in orange) from experiments done without CHX pretreatment (labeled in purple), confirming the generality of the conclusion in (B). Darker labels of each color correspond to those samples compared in (B). **(D)** Spearman rank correlations of codon identities’ A-site occupancies with the inverse of tRNA adaption index (tAI) in different experiments. A positive correlation represents translation dynamics in which codons decoded by less abundant tRNAs take longer to translate, while a negative correlation implies that codons decoded by less abundant tR-NAs are counter-intuitively faster to translate. Experiments using the standard CHX pretreatment protocol report a negative correlation, experiments done without CHX pretreatment report a positive correlation, and three sets of experiments across a gradient of CHX concentrations produced by Gerashchenko[13] each interpolate between these two phenotypes.

We compared the mean relative A-site enrichments for all 61 non-stop codons between the original CHX-pretreatment data of Ingolia et al. [21], an experiment we performed using the same CHX-pretreatment protocol, and data from experiments without CHX pretreatment by Gerashchenko [13] and by Weinberg et al. [45] (Figure 1B). A-site occupancies are strongly positively correlated between experiments that use CHX pretreatment (upper left panel) and between experiments that do not (lower right panel). The two sets of values reproducibly reported by each experimental protocol are inconsistent with each other, however, with a moderate negative correlation between them (upper right panel). To test the generality of these comparisons, we computed Pearson correlations between the A-site occupancies in representative experiments from many different studies in yeast and performed unsupervised hierarchical clustering on the resulting matrix of correlation values (Figure 1C). Experiments with and without CHX pretreatment separate into two distinct clusters, confirming that the two experimental conditions produce two reproducible but different pictures of translation dynamics.

Because the codon located in the A-site is not expected to be the only determinant of how long a ribosome spends at each position, we also calculated the influence of the codon located in the P-or E-site of a ribosome on measured ribosome density in each experiment. To do this, for each codon identity, we computed the average relative enrichment of ribosomes whose A-site was one position (P-site, Figure S2A) or two positions (E-site, Figure S3A) downstream of the codon identity, rather than directly over the codon identity. Clustering the same set of experiments by the correlations between their P-site occupancy values for all 61 codons recapitulates the same strong separation of CHX experiments from no-CHX experiments produced by the A-site occupancies (Figure S2B, C). E-site occupancy values exhibit less dynamic range from codon to codon under either experimental condition compared to the A-or P-sites (Figure S3B) but still generally separate the two conditions from each other (Figure S3C).

### CHX-induced changes in ribosomal A-and P-site enrichments are concentration dependent

The fact that tRNA binding site enrichment values from experiments with and without CHX pretreatment separate into two clusters represents two incompatible claims about how long ribosomes spend translating each codon identity. To test how well each of these two apparent phenotypes agreed with intuitive expectations about elongation times, for each experiment, we computed the Spearman rank correlation between each codon identity’s mean relative A-site enrichment and the inverse of its tRNA adaptation index (tAI) [10, 43]. The tAI of each codon identity is the weighted sum of the genomic copy numbers of the different tRNA genes that can decode the codon, with empirically determined weights penalizing wobble base pairings. This calculation quantifies the expectation that tRNAs expressed at lower abundances or that involve non-standard base pairing in their codon-anticodon interaction should require longer to translate. Consistent with previous reports [36, 5, 49, 2], all CHX experiments report weak to moderate negative correlations (Figure 1D, orange labels), representing apparent translation dynamics in which less abundant tRNAs are actually translated faster. Experiments without CHX, on the other hand, report positive correlations of varying magnitude (Figure 1D, 0x Gerashchenko NAR points and purple labels). Experiments by Pop [34], Lareau [27], Nedialkova [30], Guydosh [17] and Gardin [12] produce weak to moderate correlations, but experiments by Gerashchenko [13], Jan [23], Williams [46], Weinberg [45], and Young [48] produce fairly strong and highly statistically significant correlations.

Serendipitously, a series of experiments by Gerashchenko [13] performed to measure the effect of CHX concentration on the observed ramp in ribosome density at the 5’ end of coding sequences provide a way to confirm that CHX is directly responsible for these contradictory results. Gerashchenko produced datasets using pretreatment with a gradient of seven different CHX concentrations (0x, 1/64x, 1/16x, 1/4x, 1x, 8x, and 100x, expressed in multiples of the original protocol’s concentration of 100 *µ*g/ml) for two different cellular conditions (unstressed and oxidatively stressed cells), and using two different concentrations (0x and 1x) for heat shocked cells. Intriguingly, the rank correlation of A-site enrichment with 1 / tAI in these experiments moves smoothly from moderately negative with the highest CHX concentration to strongly positive with no CHX across each set of samples (Figure 1D), with only one sample (1/16x unstressed) deviating from perfect monotonicity. This concentration-dependence is a strong confirmation that CHX systematically shifts statistical properties of where ribosomes are measured.

To further explore the effect of CHX on A-site occupancies, we plotted the movement of the mean relative A-site enrichment of each codon identity across the concentration gradient (Figure 2 shows data from the oxidatively stressed set of samples; Figure S4 shows data from all three sets). Strikingly, a set of the codons with the highest enrichments (that is, the codons that are slowest to translate) when there is no CHX undergo consistent, gradual depletion with increasing concentration until they become among the fastest. Two prominent examples are CGA and CGG, codons encoding arginine. Mean relative enrichment at CGA codons is approximately four with no CHX, but this steadily decreases to a final value of less than 1/2 at the highest CHX concentration. CGA is translated by a tRNA identity with a moderate genomic copy number. However, its first anti-codon nucleotide is postranscriptionally modified to an inosine, making it the only codon in yeast that is decoded exclusively by an I-A wobble pairing [32]. Several studies have demonstrated that this leads to substantial translational pausing at occurrences of CGAs, particularly at CGA-CGA dicodons [28, 47, 31]. CGG, which is decoded by a tRNA with only a single genomic copy and therefore also expected to be slowly translated, undergoes an even larger shift from apparently slow with no CHX to apparently fast with high CHX concentration. For both codons, a CHX-concentration-dependent (and therefore almost certainly CHX-mediated) mechanism drives measured translation speeds away from their intuitively expected values.

**Figure 2:**
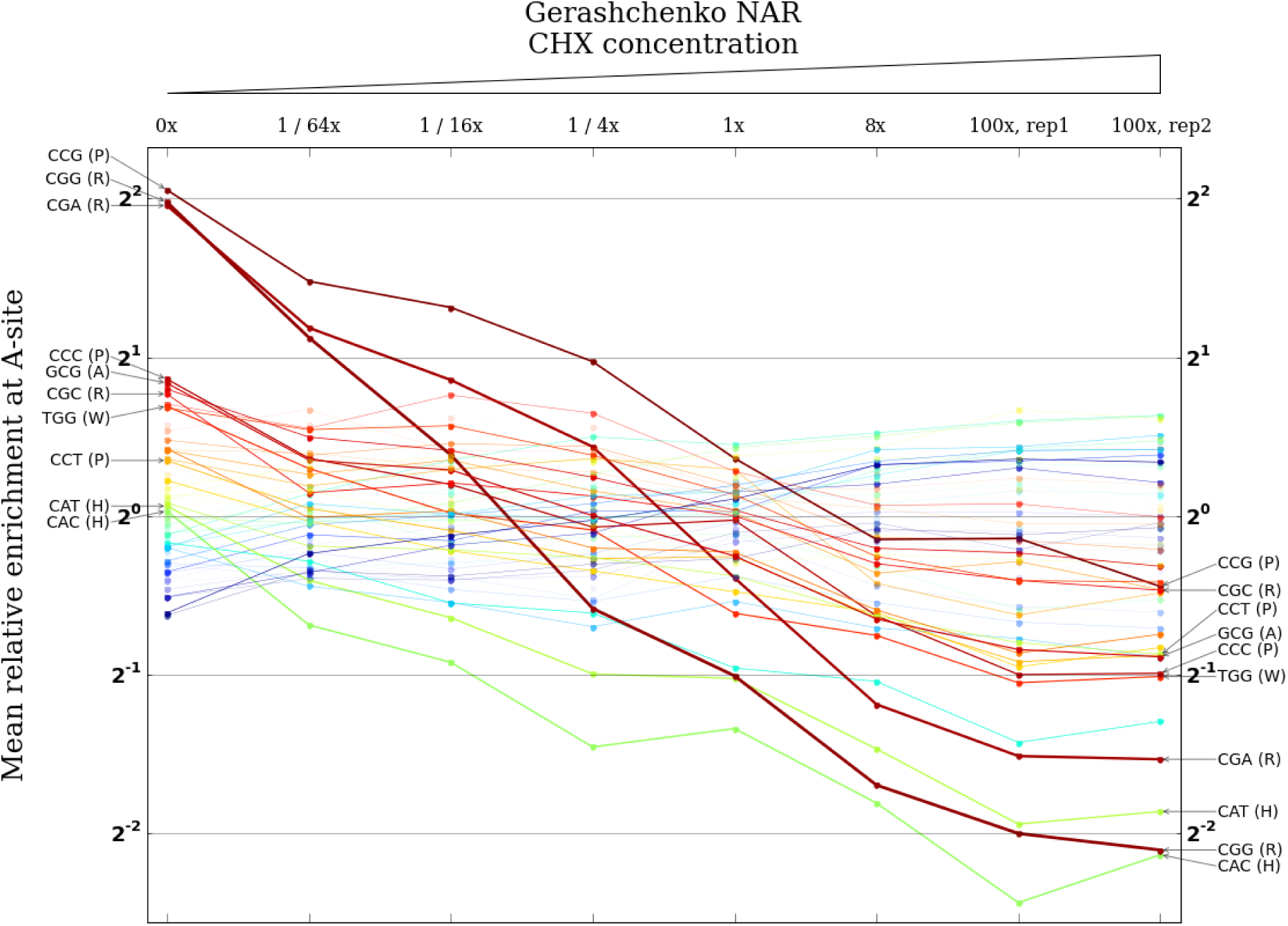
CHX pretreatment affects A-site occupancies in a coherent concentration-dependent manner. Columns correspond to a series of experiments by Gerashchenko [13] using a gradient of concentrations of CHX, starting from no CHX on the far left and increasing to 100 times the standard concentration on the far right. Each column plots the measured A-site occupancies of all 61 non-stop codons for that concentration on a log scale. The width of the line connecting each codon identity across concentrations is scaled by the codon’s net change from no CHX to 100x CHX. The ten codon identities with the largest net changes are labeled. Most notably, the codon identities with the highest enrichments in the experiment with no CHX undergo dramatic, concentration-dependent depletions over the course of the gradient.

We also examined changes in occupancy at the P-and E-sites across the concentration gradient (Figure S4). A smaller number of codons undergo substantial changes in mean relative P-site enrichment, with the dominant effect being a dramatic reduction in CGA enrichment with increasing CHX concentration. Compared to the A-and P-sites, there is less concentration-linked change in occupancy at the E-site.

### Experiments using CHX exhibit consistent patterns in ribosome density downstream of different codon identities

Although there is no *a priori* reason to expect a codon to have any impact on the translation speed of a ribosome whose A-site is more than a few positions away from it, it is straightforward to measure the average ribosome density at any particular offset upstream or downstream of a given codon identity. To do this, relative enrichments of footprint reads with their A-site at each codon position are computed as above. The relative enrichments values at all codon positions located exactly the offset of interest away from an occurrence of the codon identity of interest are then averaged (Figure 3A). We computed mean enrichment values for a wide range of offsets around each codon identity in data from our (CHX pretreatment) experiment. The A-, P-, and E-site occupancies above are the special cases of offsets of 0, +1 and +2 downstream, respectively. Although we did not expect mean enrichments to deviate substantially from one at offsets that are far removed from the tRNA binding sites, we were surprised to find prominent peaks and dips in enrichments downstream of many codon identities (Figure 3B, S5).

**Figure 3:**
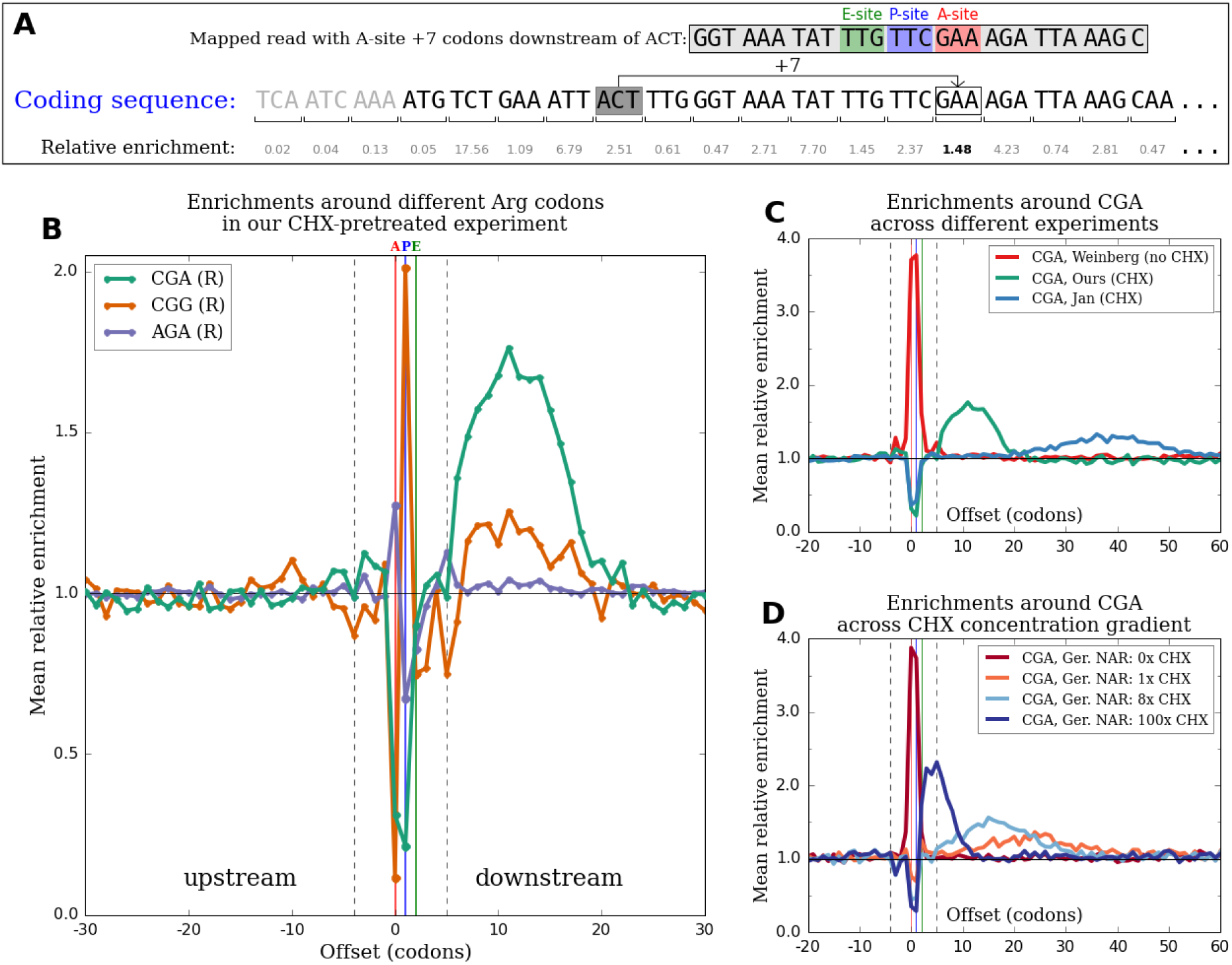
CHX pretreatment produces patterns in ribosome density downstream of different codon identities. **(A)** To measure how frequently ribosomes are observed with their A-site positioned at a particular offset upstream or downstream of a given codon identity, e.g. 7 codons downstream of an ACT (like the boxed footprint sequence on the top line), relative enrichments at each position are first calculated as in Figure 1A. The enrichment values at all positions located exactly 7 codons downstream of an ACT (such as the bolded value) across all coding sequences are then averaged. **(B)** Profiles of mean relative enrichments at a range of offsets around three arginine codons in our CHX-pretreatment experiment. Dashed vertical lines mark the boundaries of a canonical 28 nt footprint, and red, blue, and green vertical lines (corresponding to offsets of 0, +1, and +2) mark the A-, P-, and E-sites. Unexpected peaks of enrichment at downstream offsets outside of the dashed lines are observed. The magnitudes of peaks vary substantially between different codon identities encoding the same amino acid, but the horizontal extents of peaks are roughly the same across all codon identities. **(C)** Profiles of mean relative enrichments around a single codon identity (CGA) in experiments from different studies. There is no downstream peak in the no-CHX-pretreatment experiment of Weinberg et al. (red). Peaks are centered at different offsets in CHX-pretreatment experiments by different groups (green and blue) and become broader and lower when located farther downstream. **(D)** Profiles of mean relative enrichments around a single codon identity (CGA) in experiments by Gerashchenko[13] using different concentrations of CHX. With no CHX (red), there is a strong enrichment at the A-and P-sites and no downstream peak. With CHX (all other colors), there are depletions at the A-and P-sites and downstream peaks that become closer, narrower, and higher with increasing concentration.

After observing these peaks in our data, we examined data from many other experiments in yeast for evidence of similar peaks. Peaks are ubiquitous in data from experiments using CHX pretreatment (Figure 3C and D, S6), but are almost entirely absent in data from experiments that do not use CHX pretreatment (Figure 3C and D, S7, but see discussion of Pop et al. data below). For data from a particular experiment, the peaks corresponding to different codon identities occupy roughly the same range of offsets downstream (Figure 3B, S5), but across experiments carried out by different groups, the locations and shapes of the set of peaks change considerably (Figure 3C, S6). The centers of peaks vary from as close as ∼7 codons downstream in data from Nedialkova [30] to as far away as ∼50 codons downstream in data from McManus [29], with other CHX experiments densely populating the range of offsets between these observed extremes. Peaks become broader in width and smaller in maximum magnitude the farther downstream they are located.

To test if CHX treatment had a concentration-dependent effect on the locations and shapes of these peaks, we again turned to data from the CHX concentration gradient experiments of Gerashchenko. Figure 3D shows enrichment profiles downstream of CGA in one series of samples; Figure S8 shows profiles in all three series. Peaks are absent in the samples with no CHX and minimal in the samples with concentrations below 1/4x the standard concentration (with the notable exception of unstressed 1/16x, which is also a clear outlier in Figure 1D and Figure S4). For samples with concentrations greater than or equal to 1x, for which clear peaks are observed, peaks are located less far downstream and become narrower and taller as CHX concentration increases (Figure 3D).

Other studies have hypothesized that interactions between recently incorporated amino acids and the ribosome exit tunnel lead to slower ribosome movement downstream of occurrences of certain amino acids[5, 6]. There are two lines of evidence that the downstream peaks observed here are not simply the result of these effects. First, within a single sample, the magnitudes of peaks vary substantially between different codons encoding the same amino acid. As examples, the peak downstream of CGA in our experiment is substantially higher than the peaks downstream of other codons encoding arginine (Figure 3B), and GCG is the only codon encoding alanine for which there is an appreciable downstream peak (Figure S5). If these peaks were caused by interactions involving an amino acid in the nascent polypeptide chain, they should be agnostic to the codon identity used to encode the amino acid. Second, the facts that the locations of peaks change in response to changes in CHX concentration and that peaks disappear in the absence of CHX strongly suggest that the peaks are a consequence of CHX pretreatment rather than a genuine feature of translation.

### Disrupting steady-state elongation rates causes downstream peaks in analytical and simulation models

Having observed large shifts in tRNA binding site occupancies between experiments with and without CHX pretreatment and the appearance of downstream peaks in CHX experiments, we sought a model for how CHX treatment disrupts the measured positions of ribosomes that could parsimoniously explain both phenomena. To test potential models, we developed a simulation of the movement of ribosomes along yeast coding sequences; see supplement for simulation details. By incorporating different possible effects of the introduction of CHX into these simulations, we could evaluate the ability of different models to explain the observed features of the experimental data.

A natural first hypothesis is that each ribosome waits an exponentially distributed amount of time until a CHX molecule diffuses into the ribosome’s E-site and irreversibly arrests it, with shorter average waiting times when increased CHX concentration is used. If ribosomes continue to spend the same relative amounts of time on each codon while waiting for CHX to arrive, however, the position of each ribosome at the random instant of CHX arrival samples from the same steady state distribution that ribosomes occupied before CHX was introduced. We confirmed by simulation that this potential mechanism produces neither downstream peaks nor substantial changes in A-site occupancies.

We therefore considered the unexpected hypothesis that ribosomes continue to advance from codon to codon after beginning to interact with CHX, but that the relative elongation rate of each codon during this continued elongation is substantially changed from its value before CHX treatment. To model this possible behavior, we simulated translation with the relative elongation time of each codon set to its mean relative A-site enrichment as measured in a no-CHX-pretreatment experiment until steady state was reached. We then switched the relative elongation time of each codon to its A-site enrichment as measured in a CHX-pretreatment experiment and allowed translation to proceed under these new elongation rates for a short period of time. At the end of this short period of continued elongation, we recorded the positions of all ribosomes and processed the simulated ribosome footprints identically to the real experimental datasets. Interestingly, the resulting simulated enrichment profiles around different codons qualitatively reproduce both major phenomena observed in data from CHX experiments (Figure 4A).

**Figure 4:**
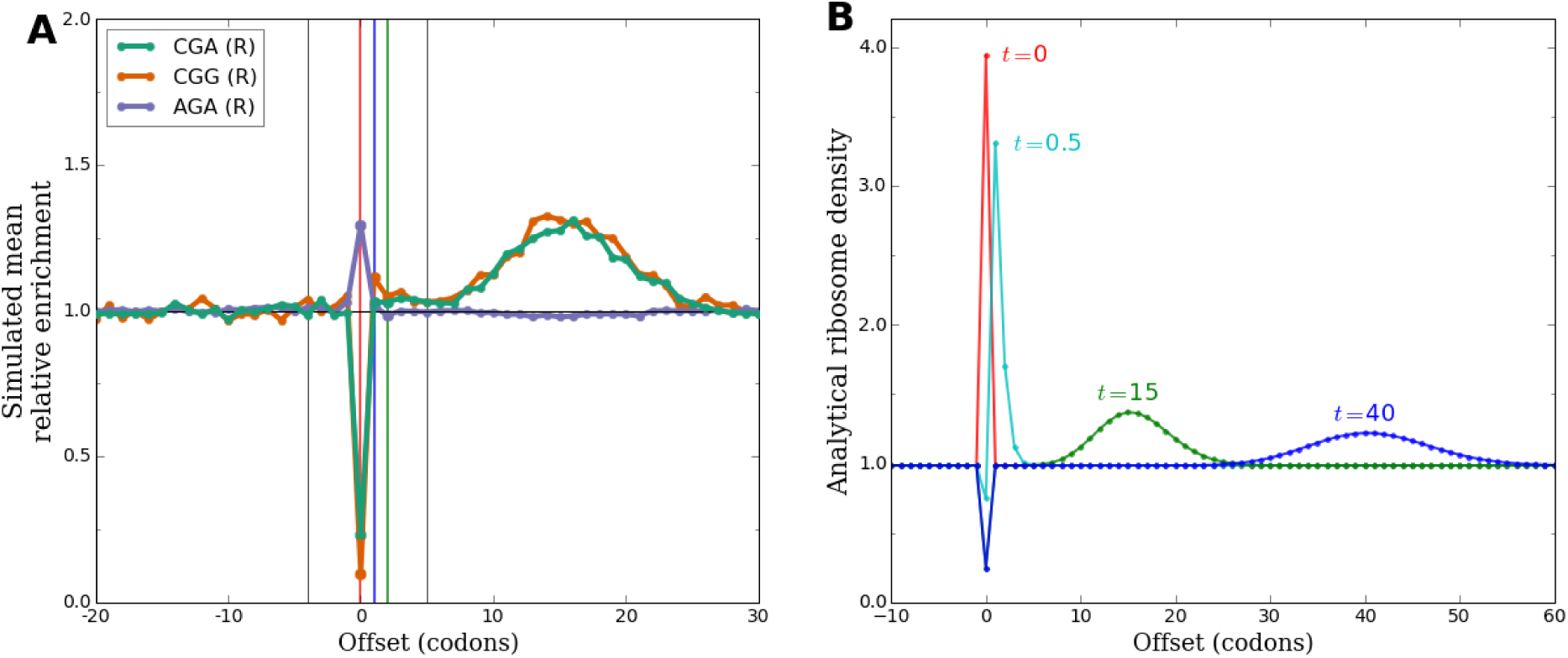
A change in the relative elongation rates of codons produces downstream waves in simulation and analytical models. **(A)** In a simulation of the translation of yeast coding sequences, the average relative elongation time of each codon identity was changed from the codon’s A-site enrichment in a no-CHX experiment to its A-site enrichment in a CHX experiment. Elongation was allowed to proceed for a short time after this change, then enrichments in the positions of ribosomes were analyzed as in Figure 3B. The resulting profiles of simulated mean enrichments qualitatively reproduce the downstream peaks in data from experiments using CHX. **(B)** A model of the translation of a hypothetical coding sequence consisting of a single slow codon surrounded by long stretches of identically faster codons on either side was analyzed. At *t* = 0 (arbitrary units), the relative speed of the slow codon was changed to be faster than its surroundings. Ribosome density at offsets around the formerly-slow codon is plotted at several time points after this change. Immediately after the change, there is a temporary excess of ribosomes positioned at the formerly-slow codon relative to the eventual steady state of the new dynamics. As these excess ribosomes advance along the coding sequence, a transient wave of increased ribosome density moves downstream and spreads out over time.

To better understand why changing relative elongation rates shortly before measuring ribosome positions produces these patterns, we constructed a simple analytical model of the translation of many copies of a particular coding sequence. In this model, ribosomes wait an exponentially distributed amount of time at each codon position before moving on to the next, with the mean of this exponential distribution depending only on the codon in the A-site of the ribosome. This implies that the steady state ribosome density at each position is proportional to the mean elongation time of the position; see supplement for details.

We considered a hypothetical coding sequence consisting of one codon that is translated slowly surrounded on either side by many identical copies of a codon that is translated faster. We computed the density of ribosomes across this coding sequence at steady state under these elongation dynamics, producing the expected excess in density at the slow codon (Figure 4B, red). Then, at *t* = 0 (arbitrary units), we changed the relative elongation rates of the two codon identities so that the previously slow codon was now faster than its surroundings. We analyzed the evolution of ribosome density across the coding sequence over time following this change. After a sufficiently long time, the system will have reached the steady state of the new dynamics, in which ribosome density is lower at the now-faster codon than its uniform level at all of the surrounding codons. Immediately after the rates are changed, however, ribosomes are still distributed at the steady state densities implied by the old dynamics and are therefore out of equilibrium under the new dynamics. There is a temporary excess of ribosomes at the formerly-slow codon, and the process of relaxing from the old steady state to the new steady state manifests as these excess ribosomes advancing along the coding sequence over time (Figure 4B). Stochastic variation in the wait times of each individual ribosome at each subsequent codon position causes the excess to gradually spread out as it advances. Hypothetical measurements of the positions of all ribosomes at a series of increasing times after the change to the new dynamics would therefore produces patterns that look like an advancing wave of enrichment, as is seen around e.g. several arginine codons in real (Figure 4A) data.

We also considered a hypothetical coding sequence in which a single special codon undergoes an increase, rather than decrease, in relative elongation time compared to stretches of identical codons on either side (Figure S9). In this case, the time period immediately following the change in dynamics is spent filling the formerly-faster codon position up to its newly increased steady state density. During this time, there are temporarily fewer ribosomes being promoted onward to downstream positions than there were before the change. This results in a transient wave of depletion, rather than enrichment, that advances away from the formerly-faster codon position and spreads out over time (Figure S9B). This is qualitatively consistent with the profile of depletions downstream of e.g. two isoleucine codons in real (Figure S5) and simulated (Figure S9A) data.

### Magnitudes and locations of downstream peaks are quantitatively consistent with predictions made by wave hypothesis

The hypothesis that changes in measured tRNA binding site occupancies and the appearance of downstream peaks are both caused by continued elongation with disrupted dynamics in the presence of CHX makes a testable prediction about the quantitative link between these two phenomena. If downstream peaks are transient waves moving downstream after a change in the relative amounts of time ribosomes spend positioned over each codon, the total CHX-induced excess or deficit in enrichment downstream of each codon identity should exactly offset the total CHX-induced change in enrichments at the tRNA binding sites. To test whether experimental data agreed with this prediction, we analyzed several matched pair of experiments performed with and without CHX by Jan [23] (Figure 5), Williams [46], and Gerashchenko [13] (Figure S11). For each codon identity, we compared the sum of the differences in enrichment between the experiments at the A-, P-, and E-sites (green area in insets) to the sum of the difference in enrichment across the range of downstream offsets occupied by the putative waves (red area in insets). In all matched pairs of experiments, the area of each codon’s downstream peak is accurately predicted by its tRNA binding site changes (*r*^2^ = 0.85 to 0.93, slope of best fit line *β* = −1.00 to −1.20). Insets in Figure 5 and Figure S10 plot the changes in tRNA binding site enrichments and downstream wave areas for four different codons to demonstrate the full range of agreement between the two phenomena. CGA, which undergoes comparably large decreases in enrichment at both the A-and P-sites, produces a downstream wave with approximately twice the area of CCG, which undergoes a large decrease in enrichment at the A-site but not the P-site. Codons with similar enrichments at all three tRNA binding sites between the two experiments, such as ACT, produce no appreciable downstream waves, while several codons that undergo modest increases in enrichment at the binding sites, such as TTG, produce proportionally modest net deficits of enrichment downstream. This close correspondence strongly suggests that the downstream peaks are in fact transient waves, and therefore that tRNA binding site enrichments in CHX experiments do not reflect natural translation dynamics.

**Figure 5:**
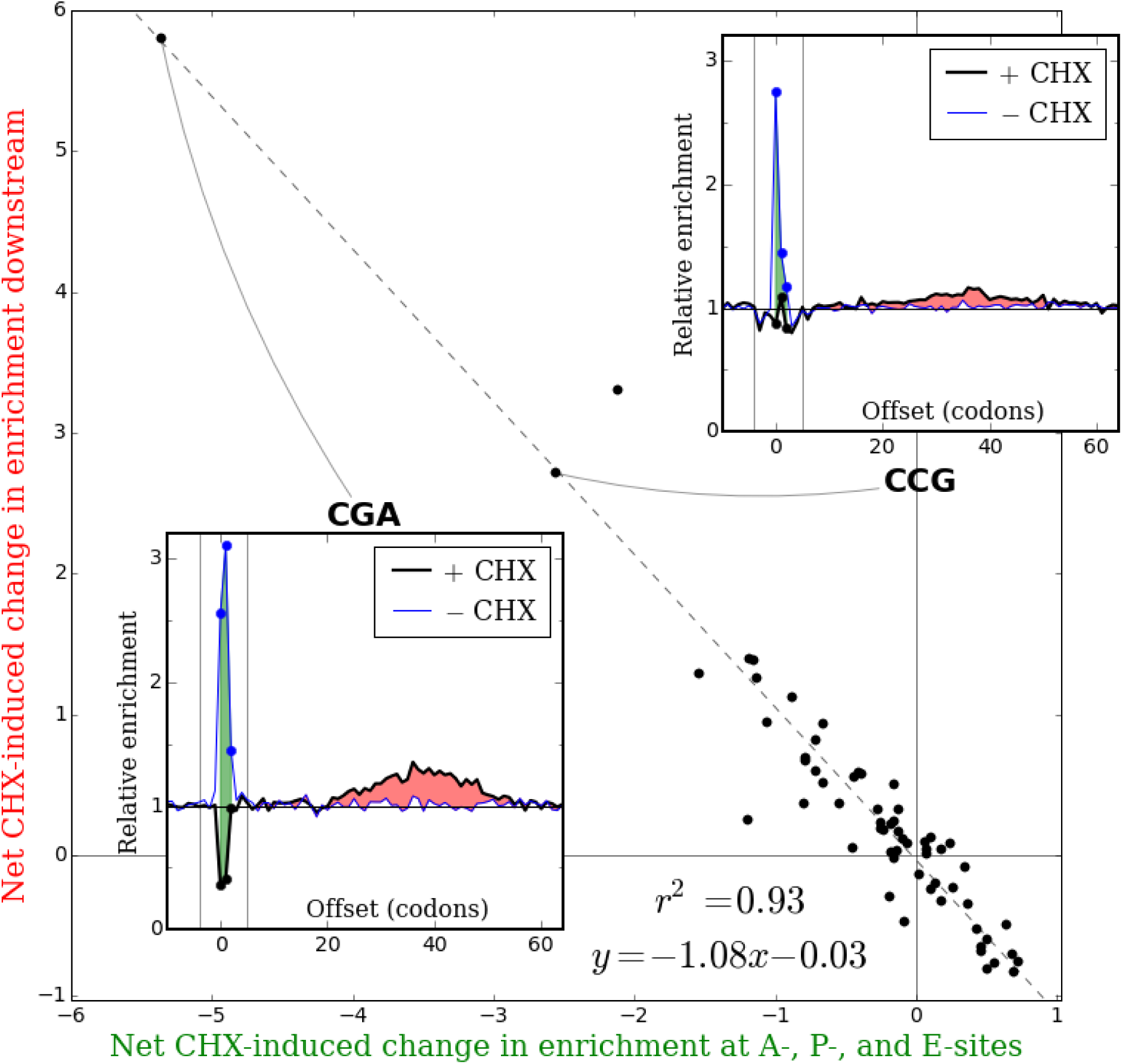
Changes in tRNA binding site enrichments between a pair of experiments with and without CHX are matched by areas of downstream waves in the CHX experiment. For a pair of experiments with and without CHX by Jan [23], the sum of each codon identity’s changes in mean relative enrichment at the A-, P-, and E-sites between the two experiments (green area in insets) is plotted against the total excess or deficit of enrichment in the CHX experiment from 6 to 65 codons downstream (red area in insets). The area of each codon identity’s downstream peak is strongly predicted by changes in enrichment at the tRNA binding sites, consistent with the hypothesis that downstream peaks are transient waves caused by continued elongation with disrupted dynamics in the presence of CHX.

A model of continuing elongation after CHX has begun interacting with ribosomes also predicts that CHX pretreatment for increasing amounts of time under otherwise identical conditions should produce waves that have advanced proportionally farther downstream. To test this prediction, we examined mean relative enrichments around CGA in the full set of experiments from Jan et al. [23], which involved a variety of different combinations of CHX pretreatment duration and temperature. Strikingly, when pretreatment was performed at 30°C, downstream peaks after 9 minutes of pretreatment are centered slightly more than twice as far downstream as those after 4 minutes of pretreatment (Figure 6), further supporting the hypothesis that elongation continues to occur at a slow but steady rate over the course of CHX pretreatment.

**Figure 6:**
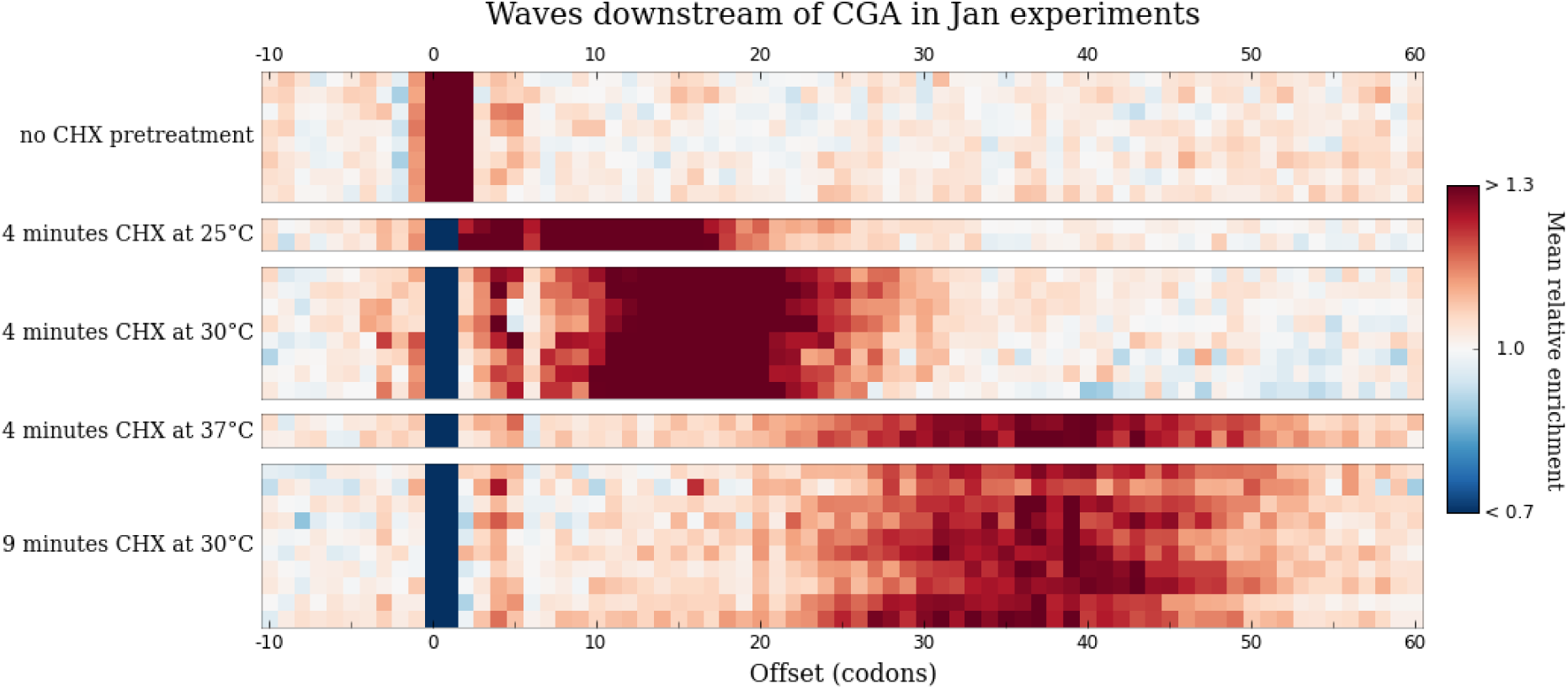
Waves move proportionally farther downstream with increasing CHX pretreatment time. Each heatmap row shows mean relative enrichments around CGA in a different experiment from Jan et al. [23], with columns corresponding to different offsets. Experiments are separated by CHX pretreatment conditions: either no pretreatment (top group) or the duration and temperature of pretreatment (all other groups). At 30°C, pretreatment for 9 minutes (bottom group) produces waves that have moved slightly more than twice as far downstream as those produced by pretreatment for 4 minutes (middle group).

### Disrupted elongation in the presence of CHX explains counter-intuitive results in CHX experiments

Under this model of continuing elongation, the areas of downstream waves can be used to recover indirect information about the history of translation dynamics in each CHX-pretreatment experiment before these dynamics were disrupted by CHX. Specifically, we can estimate what the sum of the enrichments at the A-, P-, and E-sites was for each codon before the introduction of CHX by adding the net area of the wave that moved downstream during elongation in the presence of CHX back to the sum of the enrichments that remain at the binding sites. We will call this quantity the corrected aggregate enrichment of each codon. It can be interpreted as the average relative amount of time that a ribosome took to decode each occurrence of a codon before CHX was introduced, from when the codon was presented in the A-site to when it left the E-site. While we would prefer to recover how long each codon spent in each individual tRNA binding site in these experiments, this single-codon-resolution information has been irreversibly lost. As the CGA and CCG insets in Figure 5 demonstrate, changes in enrichment at the A-site or at the P-site result in downstream waves that occupy the same large range of downstream offsets, so the area in each wave cannot be unambiguously assigned back to a particular tRNA binding site.

To test if codons decoded by less abundant or wobble-paired tRNAs tended to be translated more slowly than more abundant tRNAs in CHX experiments before the introduction of CHX, we computed the Spearman rank correlation between the corrected aggregate enrichment of each codon identity and 1 / tAI. Corrected aggregate enrichment correlates positively with 1 / tAI for every CHX experiment analyzed (Figure 7, purple dots), recovering an intuitively expected signature of translation dynamics that is absent in CHX experiments if the total elongation time is estimated by the sum of the tRNA binding sites enrichments alone (Figure 7, green dots).

**Figure 7:**
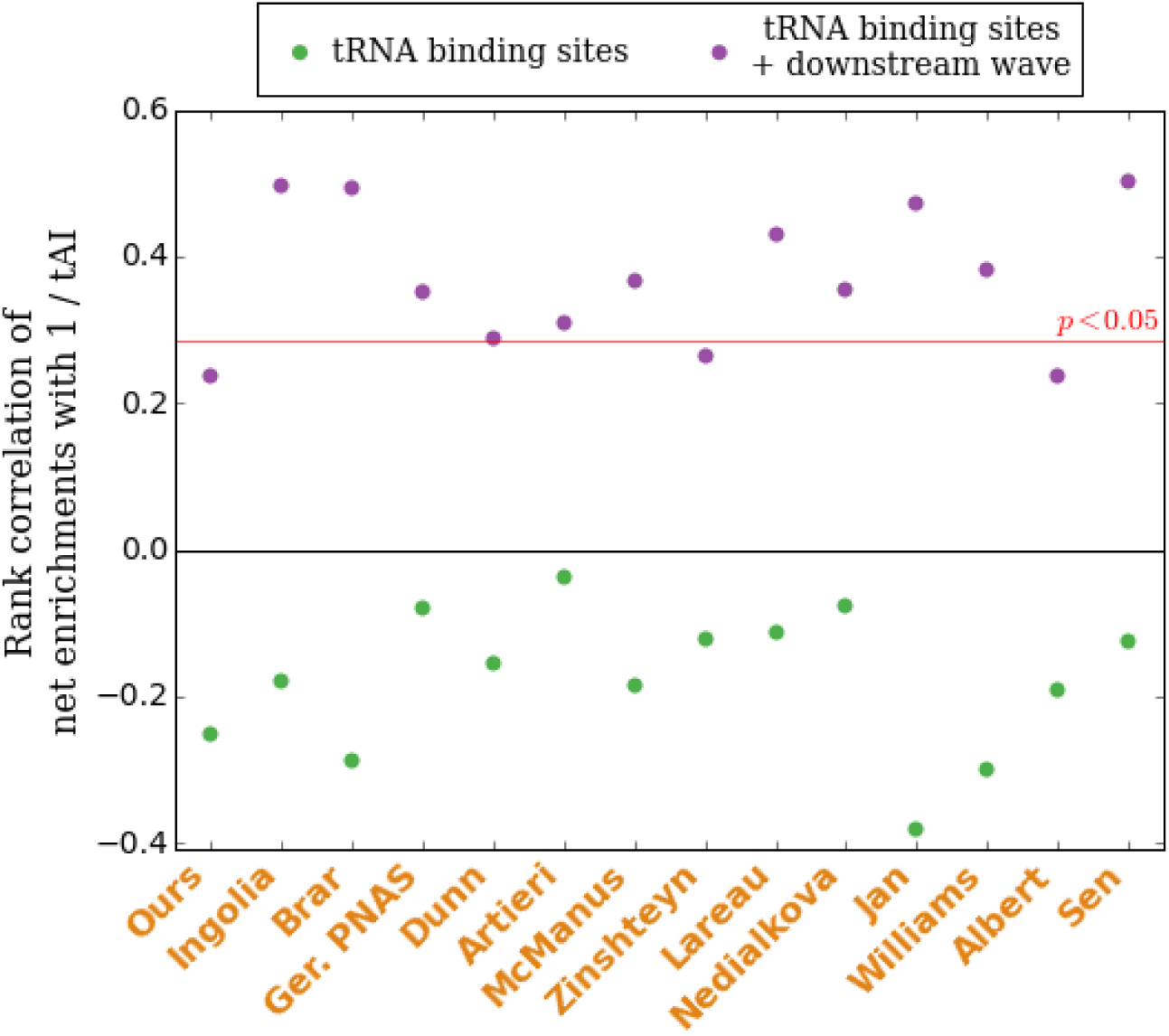
Downstream waves recover positive correlations of estimated elongation times with 1 / tAI. In experiments using CHX pretreatment, the combined enrichment of each codon identity at the A-, P-, and E-sites before the introduction of CHX can be estimated by adding the net area of the codon’s downstream wave back to the total remaining enrichments at the three tRNA binding sites. This sum correlates positively with 1 / tAI in all CHX experiments (purple dots), recovering the positive correlations counterintutively absent at the tRNA binding sites alone in these experiments (green dots). Positive correlations are statistically significant (*p*< 0.05, one-tailed) in most experiments. This suggests that non-optimal codons were being translated less quickly than optimal codons in CHX experiments before the introduction of CHX disrupted these dynamics.

Continued elongation with disrupted dynamics after the introduction of CHX also offers a potential explanation for counterintuitive results in a set of experiments by Zinshteyn et al. [49]. Zinshteyn performed ribosome profiling on yeast strains that lacked different genes required to post-transcriptionally add mcm^5^s^2^ groups to a uridine in the anticodons of tRNAs that decode codons ending in AA and AG. These anticodon modifications are thought to enhance codon-anticodon recognition and speed up translation of these codons [37]. Surprisingly, Zinshteyn found that measured changes in tRNA binding site occupancies between deletion strains and the wild type were much smaller than expected given the phenotypic consequences of lacking these modifications. These experiments followed the standard CHX pretreatment protocol, however, and we observe clear downstream waves in enrichment in all of them (Figure S12). According to our model, therefore, tRNA binding site occupancy levels in these experiments reflect properties of elongation in the presence of CHX rather than of *in vivo* dynamics.

To test if CHX-disrupted elongation was masking the true impact of the absence of anticodon modification in these experiments, we compared the profiles of mean enrichment around all codon identities decoded by the modification-deficit tRNA species between the deletion strains and the wild type. Intriguingly, the profiles of mean enrichment around AAA showed consistently increased downstream wave areas in all of the deletion strains compared to wild type (Figure 8A). To quantify this increase, we computed the corrected aggregate enrichment of each codon identity as above by adding the downstream wave area to the sum of the binding site enrichments. We then computed the change in corrected aggregate enrichment for each codon identity between each of the deletion strains and the wild type (Figure 8B, S13). In each of the deletion strains, but not in a replicate of the wild type, AAA undergoes a dramatically larger increase in aggregate tRNA binding site enrichment when corrected to include downstream wave area (purple) than if wave area is not included (green). This argues that AAA does in fact take substantially longer to decode *in vivo* in cells lacking the ability to modify its tRNA, but that most of this difference disappears during continued elongation in the presence of CHX.

**Figure 8:**
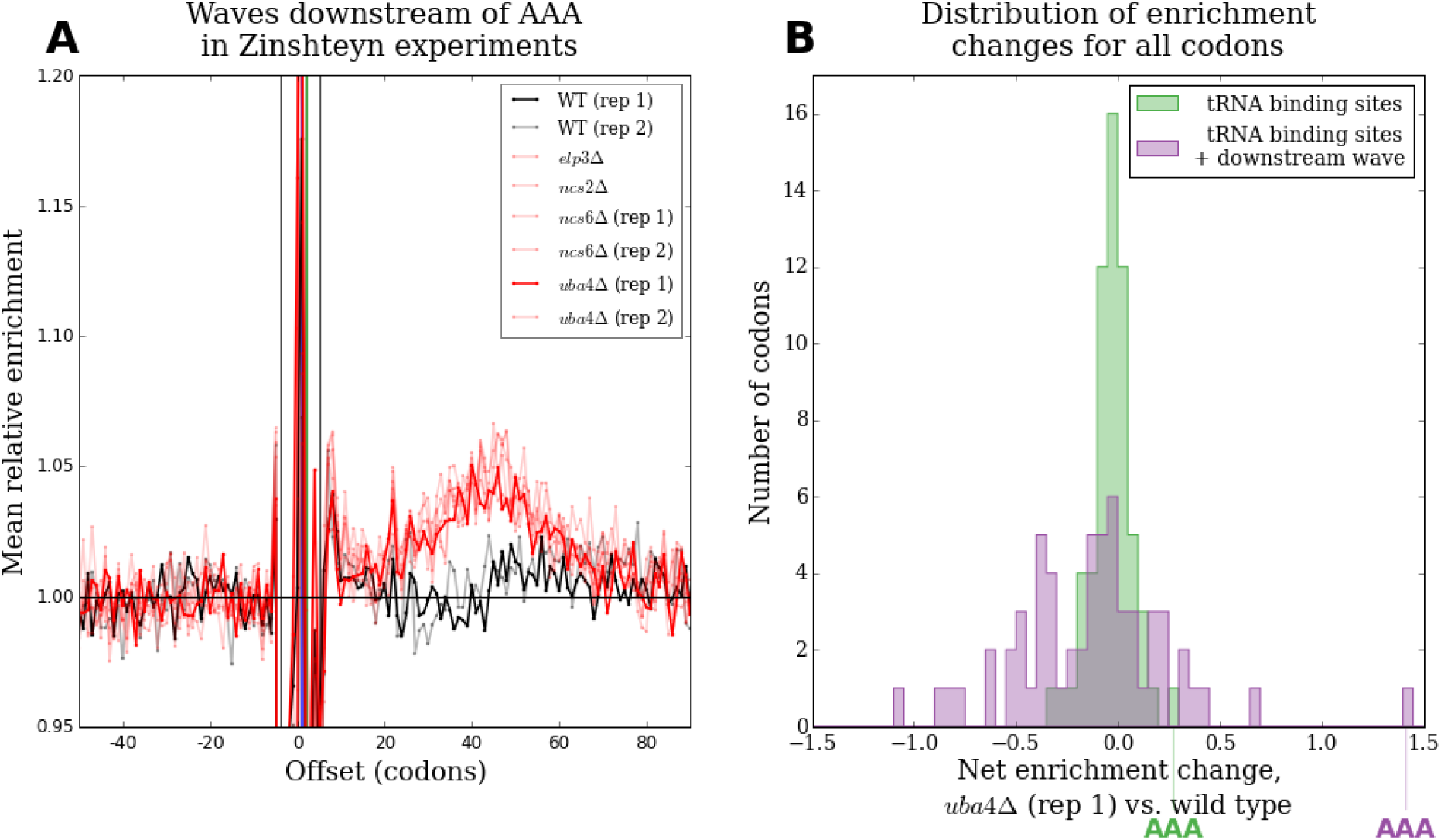
Downstream waves recover the expected effects of lacking tRNA modifications. **(A)** Profiles of mean relative enrichments around AAA in two wild type experiments (black lines) and six experiments with different components of the mcm^5^s^2^U pathway deleted (red lines) from Zinshteyn [49]. All mcm^5^s^2^U deletion strains produce clearly increased waves downstream of AAA compared to wild type. Darker lines correspond to the experiments compared in (B). **(B)** Histograms of the net change in enrichment for each codon identity between *uba*4Δ and wild type at the A-, P-, and E-sites (green) or at the A-P, and E-sites plus 7 to 90 codons downstream (purple). AAA shows a modest increase in net enrichment at the tRNA binding sites, but a dramatically larger increase in net enrichment if the area of downstream waves is also taken into account. This suggests that AAA does take substantially longer to decode *in vivo* in *uba*4Δ than in wild type, but that most of this difference disappears during continued elongation in the presence of CHX.

## Discussion

We have seen that ribosome profiling experiments in yeast performed with and without CHX pretreatment make incompatible claims about the average amounts of time ribosomes spend with each codon identity in their tRNA binding sites during elongation. We have reported the existence of unexpected structure in measured ribosomes density downstream of each codon identity in experiments using CHX pretreatment. The most parsimonious model to explain both of these phenomena is that elongation continues for many cycles after the introduction of CHX, but that during this continued elongation, the amount of time each ribosome takes to advance from codon to codon has a different quantitative dependence on the codons positioned in the tRNA binding sites of the ribosome than it did before the introduction of CHX.

The interpretation of experiments using CHX pretreatment to stabilize ribosomes has assumed that this stabilization occurs by a mechanism that leaves ribosomes positioned according to their steady state *in vivo* distribution, e.g. by irreversibly arresting the further elongation of each ribosome upon binding to it. The patterns we observe argue that this assumption does not hold. Instead, many cycles of continued elongation with disrupted dynamics leave ribosomes distributed in a way that does not directly reflect natural translation dynamics, although telltale signs of the pre-disruption dynamics can be indirectly discerned. In light of this, future ribosome profiling experiments aiming to directly measure the amount of time ribosomes spend at each position *in vivo* should entirely avoid the use of CHX.

### Mechanism of continued elongation in the presence of CHX

Although the exact mechanistic details of how disrupted elongation in the presence of CHX occurs remains unclear, there are several key features of observed patterns in the data and of known properties of CHX that any potential mechanism must accommodate. The first is that the disruption in dynamics is concentration-dependent. The second is that relative elongation rates in the new dynamics are still coupled to codon identities. Mean relative A-and P-site enrichments do not simply collapse towards being uniform in CHX experiments, but instead reproducibly take on a wide dynamic range of codon-specific values. The third is that absolute elongation rates must be dramatically slower in the presence of standard concentrations of CHX. Any model implying the contrary is not plausible; CHX has been successfully used as a translation inhibitor for decades. We suggest that this inhibition is accomplished by a large reduction in the rate of elongation rather than a complete halt.

A possible mechanism with all three of these properties is that CHX repeatedly binds and unbinds ribosomes, preventing advancement when bound but allowing elongation to proceed when unbound. In this model, the global rate of CHX binding to all ribosomes increases with increasing CHX concentration, leading to a decrease in the amount of time each ribosome spends unbound and therefore globally decreasing the rate of continued elongation. This accounts for the fact that downstream peaks move less far downstream in the same amount of time with increasing CHX concentration (Figure 3D). For fixed CHX concentration, there is close proportionality between the duration of CHX pretreatment and the distance peaks have moved downstream in experiments performed by the same group (Figure 6). While comparisons of experiments by different groups may be confounded by subtle differences in experimental protocols, there is broad (but not perfect) agreement between downstream peak distance and annotated pretreatment time across experiments from different studies, with longer pretreatment times (e.g. 5 minutes in McManus[29]) corresponding to peaks farther downstream and shorter pretreatment times (e.g. 1 minute in Lareau [27] and Nedialkova [30]) corresponding to closer peaks. Because the distance that peaks have moved downstream is the product of the total duration of disrupted elongation and the average rate of this elongation, the magnitude of the reduction in elongation rate can be roughly estimated. The range of peak centers with standard CHX concentration and 2 minutes of pretreatment implies absolute elongation rates during CHX treatment of 0.1 to 0.3 aa/s. Because natural elongation rates in yeast are 7 to 9 aa/s [26], this represents an approximately 20-to 90-fold reduction in the speed of elongation.

To explain the reproducible range of codon-specific elongation rates in the presence of CHX, changes in the conformation of the ribosome as a result of differences in the geometry or base-pairing interactions of the tRNAs occupying the A-and P-sites could modulate the rates of CHX binding and unbinding. Conformational changes in the ribosome as a result of codon-anticodon interactions are known to be an integral part of the elongation cycle [42]. Given the unique presence of I-A wobble pairing in the decoding of CGA codons, the outsize role that CGA plays in these phenomena suggests that base-pairing interactions could play a major role in determining CHX affinity. This offers an elegant potential explanation for the negative correlation between A-site enrichments with and without CHX: codon identities that produce unusual ribosome conformations tend to slow down elongation when tRNA binding is rate-limiting, but tend to speed up elongation when CHX disassociation is rate-limiting. In this model, the concentration-dependent interpolation between these two regimes observed in Figure 2 reflects the fact that as CHX concentration decreases, each ribosome spends an increasing fraction of time unbound by CHX and therefore elongating according to the unperturbed dynamics.

### Heterogeneity in experiments without CHX pretreatment

Heterogeneity in measured A-and P-site enrichment values between experiments that avoid CHX pretreatment presents complications for this model. Except for two experiments in the set performed by Guydosh [17], however, all such experiments still include CHX in the lysis buffer into which cells are harvested. Under some conditions, the same continued elongation with disrupted dynamics that occurs during CHX pretreatment could also occur during the harvesting process once ribosomes have been exposed to CHX in the lysis buffer. The A-and P-site occupancies in the non-pretreated experiments by Pop, Lareau, Gardin, Nedialkova, and the unstressed experiment by Gerashchenko can be interpreted as an intermediate phenotype halfway in between the two tighter clusters consisting of CHX-pretreatment experiments and of the non-pretreated experiments by Weinberg, Jan, Williams, Guydosh, Young, and the other two experiments by Gerashchenko (Figure 1B), potentially reflecting a small amount of CHX-mediated elongation in these intermediate experiments. Consistent with this interpretation, enrichment profiles around CGA and CGG appear to be shifted slightly downstream in these intermediate-phenotype non-pretreated experiments (Figures S15, S16).

The complete set of five non-pretreated experiments produced by Pop et al.[34] are particularly heterogeneous. Three of these experiments (WT-URA footprint, AGGOE footprint, and AGG-QC_footprint) report A-site occupancies strikingly similar to values reported by CHX pretreatment experiments and less similar to the other non-pretreated experiments (Figure S14). These same three experiments also have distinct peaks located approximately 20 codon positions downstream in the enrichment profiles around each codon identity that are absent in the other two experiments from the study (Figure S16). Such extreme heterogeneity between non-pretreated experiments is difficult to account for in our model, but suggests that a wide range of different amounts of elongation after exposure to the lysis buffer are possible across different implementations of harvesting protocols.

### Reexamination of conclusions from CHX-pretreatment experiments

In light of our model, several counterintuitive results from previous ribosome profiling studies in yeast take on new interpretations. Most notably, we offer an explanation for contradictory claims about whether so-called optimal codons corresponding to more abundant tRNAs are translated more rapidly. In many organisms, optimal codons are used with greater frequency in highly expressed genes, and a large body of theoretical work assumes that increased elongation speed drives this tendency [33]. If optimal codons are decoded faster, this tendency could lead directly to increased expression by increasing the rate of production of protein from each message or by avoiding mRNA-decay pathways linked to ribosome pausing [35]. Alternatively, if translation initiation is typically rate-limiting, using faster codons reduces the amount of time ribosomes spend sequestered on highly transcribed mRNAs, freeing up ribosomes to translate other messages and leading to more efficient system-wide translation [40]. If A-site enrichments measured in CHX pretreatment experiments reflected natural translation dynamics, however, optimal codons would not be elongated more quickly, and these theories fall apart. By offering a model for why the measured positions of ribosomes in CHX experiments appear to report that non-optimal codons are the fastest to be translated, and by showing evidence that optimal codons were in fact being translated more quickly before the introduction of CHX, we enable a principled resolution to this controversy. Earlier studies in this area have hypothesized that the A-site enrichments reported by CHX experiments could reflect an optimal balance between codon usage and tRNA abundance [36], or that potential heterogeneity in elongation times at different occurrences of the same codon identity could conspire to produce these A-site enrichments [7]. By showing that the A-site occupancies fed into these models almost certainly do not represent the actual *in vivo dynamics*, our results argue against both of these conclusions.

Finally, our model offers an explanation for the small apparent impacts of several experimental attempts to modify tRNA repertoires on measured relative elongation rates. CHX-pretreatment experiments by Zinshteyn et al. [49] on mcm^5^s^2^U-pathway deletion strains show surprisingly small changes in tRNA binding site occupancies at codons decoded by the modification-deficient tRNAs. Our observation that all of the deletion strains have substantially increased waves of enrichment downstream of AAA, one such codon identity, compared to wild type suggests that binding site occupancy changes in the presence of CHX dramatically underestimate the actual *in vivo* increase in the decoding time of AAA in the deletion strains. Pop et al. [34] evaluated the impact of overexpressing, deleting, or modifying the body sequence of tRNAs and also found surprisingly small changes in the rates at which the corresponding codon identities were translated. As discussed above, the experiments of Pop et al. did not pretreat with CHX, but a subset of these experiments show both clear downstream peaks and A-site occupancies shifted towards values reported by CHX pretreatment experiments. This suggests that enough CHX-disrupted elongation occurred in these experiments during the harvesting process that the resulting A-site occupancies may not be able to measure any potential effects of the tRNA repertoire modifications. Repeating these experiments without any CHX in order to accurately sample from *in vivo* dynamics could clarify the consequences of these tRNA manipulations.

## Acknowledgments

We thank John Hawkins, Alex Stabell, Premal Shah, Joshua Plotkin, David Weinberg, and David Bartel for helpful discussions, Kim DeKeersmaecker for sequencing resources, and Christina Pop and Liana Lareau for clarifications about their experimental protocols.

